# Microbial metabolism of mannitol as a tracer for the non-invasive measurement of oral-cecal transit

**DOI:** 10.1101/2025.10.28.685202

**Authors:** Daniel E. Vieira, Emily M. Langmeyer, Marissa D. Cortopassi, Deepti Ramachandran, William B. Rubio, Nagasuryaprasad Kotikalapudi, Yancheng Zhou, Sarah N. Flier, Lynn Bry, Alexander S. Banks

**Affiliations:** Division of Endocrinology, Diabetes and Metabolism, Beth Israel Deaconess Medical Center, Harvard Medical School, Boston, MA, USA; Division of Gastroenterology, Beth Israel Deaconess Medical Center, Boston, MA, USA; Massachusetts Host-Microbiome Center, Brigham and Women’s Hospital, Harvard Medical School, Boston, MA, USA

## Abstract

The timing of digestion after a meal reveals information essential for evaluating gastrointestinal function and overall digestive health. Alterations in transit time through the GI tract can reveal abnormalities such as delayed gastric emptying and provide critical insights into nutrient absorption dynamics. This information is particularly valuable for optimizing dietary interventions, managing metabolic conditions such as diabetes, and improving personalized nutrition strategies. Methods for measuring the oral to cecal transit time (OCTT) in murine models have significant limitations. We demonstrate a non-invasive approach in freely-moving non-anesthetized mice which quantifies the microbial digestion of the non-nutritive sweetener mannitol to CO_2_. We monitor cage air for the production of ^13^CO_2_ from ^13^C-enriched mannitol using Off-Axis Integrated Cavity Output Spectroscopy integrated with multiplexes indirect calorimetry. With this approach, we find mannitol oxidation is absent in mice following commensal depletion of the microbiota. In mice with conventional microbiota, the peak ^13^C-mannitol oxidation occurs proximal to the mouse cecum, allowing the quantitation of OCTT in mice. By tracking the output of ^13^CO_2_, this method provides highly granular, real-time data. We detect delayed OCTT with the use of pharmacological transit-altering compounds loperamide, a gut restricted opioid receptor agonist and also semaglutide, a GLP-1 receptor agonist. This approach may provide more physiologically relevant results in a range of genetic, environmental, and pharmacological research models.

## Introduction

Understanding gastrointestinal transit times is essential for studying gut physiology, but current methods for measuring oral cecal transit time (OCTT) in murine models present significant limitations. Existing approaches, such as oral administration of fluorescently labeled polymers (e.g. FITC dextran) or metallic beads followed by dissection of the GI tract can offer high granularity, but may confound results due to the use or anesthesia or physiological invasiveness (1, 2). These methods, including physical examination of dye fronts within the murine GI tract, can provide a high level of accuracy but are limited by the fact that they are terminal procedures. Available non-terminal whole gut transit measurements include the Carmine red assay, which involves regular examination of animals to determine when the dye has been excreted. These examinations of the mice during the assay can drastically decrease transit time due to stress; and although automated methods have been developed to improve this assay, it still lacks the resolution of the aforementioned terminal procedures. Other non-terminal alternatives, such as Single-Photon Emission Computed Tomography (SPECT) after administration of 99mTc-DTPA-labeled activated charcoal, require anesthetization of murine models and use of short half-life radioisotopes (3, 4). These constraints present an opportunity for the development of a non-invasive technique for accurately measuring OCTT. Such a technique would balance accuracy and precision, with minimal animal stress while maintaining the ability to collect longitudinal data in real time. The ideal method should eliminate the need for terminal procedures, anesthesia, or repeated handling of animals, thereby reducing experimental bias and improving the translational relevance of murine gastrointestinal studies. Furthermore, a noninvasive method to measure gastrointestinal transit times in mice would significantly advance our ability to study motility, nutrient absorption, and the complex interactions between diet, microbiota, and host physiology as well as the impact of pharmacologic therapies.

The use of real-time stable isotope measurement is well established in both clinical and preclinical research (3-5). The gastric emptying breath test measures the oxidation of ^13^C-labeled octanoic acid (OA), a medium-length fatty acid absorbed early in the duodenum. Once absorbed, OA is metabolized in the liver, and a fraction is exhaled as ^13^CO_2_. The time course of ^13^CO_2_ production reveals information about the rate at which solid food is emptied from the stomach and reaches the duodenum. However, detection of stable isotope tracer-metabolites more distally in the GI tract is not commonplace due to the lack of appropriate reagents.

Sugar alcohols, including mannitol, sorbitol, and xylitol are used as artificial sweeteners as they produce a sweet taste sensation and increase the palatability of foods but are poorly absorbed in the human gut (6). Mannitol occurs naturally in watermelon, peaches, and olives (7, 8). As a food additive, mannitol does not promote weight gain in animal models (9). At high concentrations, orally ingested mannitol can produce a laxative effect (10), but these effects are not commonly seen in doses used for sweeteners.

Here we demonstrate ^13^C-labelled mannitol is metabolized by microbiota in the murine cecum. This conversion of ^13^C-mannitol to ^13^CO_2_ establishes a highly granular, non-invasive approach to measure OCTT in mice. This method circumvents the need for terminal procedures, significantly reducing the number of animals required for transit-time experiments, while still providing the quality and granularity of those types of procedures. This study uses new indirect calorimetry technology to detect total CO_2_ production combined with a real-time detector of ^13^C/^12^C ratio. This allows for rapid, sequential, measurement of multiple cages simultaneously. These capabilities can increase the throughput of preclinical studies while also limiting potential extraneous variables. The ability to conduct repeat studies with greater ease and consistency enhances reproducibility and robustness of findings. This advancement could represent a major step forward in experimental design, aligning scientific rigor with ethical considerations in animal research.

## Results

### Detection of gastric emptying through ^13^C-octanoic acid oxidation

We sought to validate our system using the established octanoic acid gastric emptying assay to measure transit time to the duodenum. We fed ^13^C-octanoic acid-containing food to individually housed, unanesthetized, freely moving mice with ad libitum access to food and water to measure the rate of gastric emptying. After consumption of a meal containing ^13^C-octanoic acid, rates of ^13^C-CO_2_ detection began to peak 50 minutes post-administration (**Figure 1A**). Roughly 2.5% of the consumed tracer was oxidized to exhaled ^13^CO_2_ by the end of the observation period **(Figure 1B)**. We next evaluated the timing of gastric emptying following administration of a prokinetic agent, metoclopramide (MCP), used to treat gastroparesis and as an antiemetic. In patients, MCP acts as an enteric dopamine receptor (D2) antagonist to accelerate movement of food through the stomach and proximal small intestine. We examined the effect of MCP on kinetics of ^13^C-octanoic acid oxidation. After administration of the tracer, we measured ^13^CO_2_ production for the next 200 minutes. We observed a sharp peak of ^13^CO_2_ occurring within 30 minutes, this is indicative of accelerated gastric emptying (**Figure 1D**). In this interval, similar amounts of total tracer were oxidized to ^13^CO_2_ in both control and MCP treated groups (**Figure 1E**). To quantify the rate of gastric emptying, we measured the fraction of tracer oxidized within the 200-minute experimental monitoring period. The time at which 50% of tracer was oxidized, T_50%_ was determined for mice in both groups. The T_50%_ of MCP-treated mice was significantly lower than the T_50%_ of saline-treated mice (**Figure 1F-G**). As we can effectively monitor ^13^C-octanoic acid oxidation, in a multiplexed non-invasive system we next sought to identify a tracer that would be absorbed or metabolized at a more distal site in the gastrointestinal tract.

**Figure 1:**
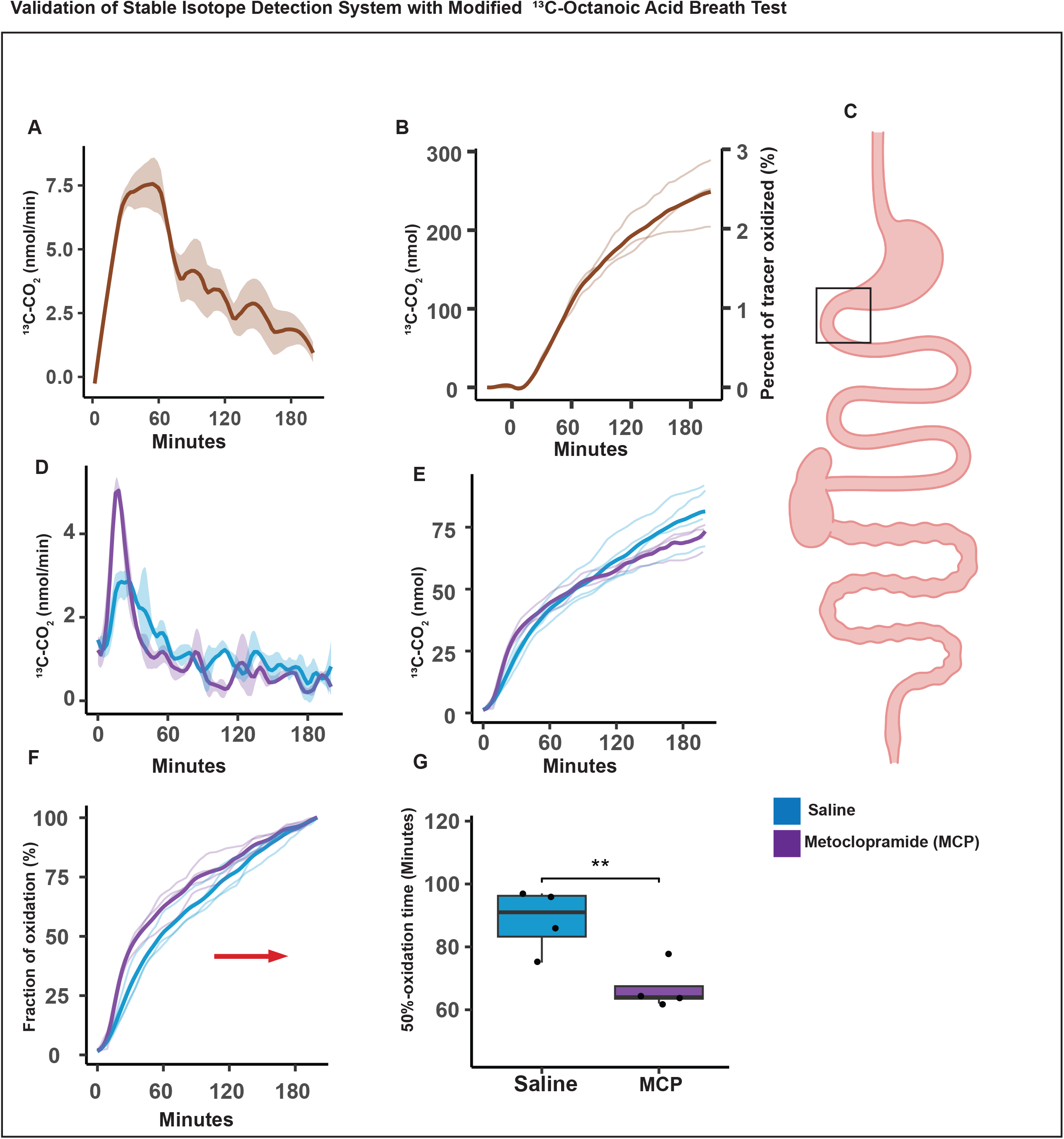
[^13^C]-octanoic acid breath test in mice. Mice voluntarily ate a meal with trace amounts of [^13^C] octanoic acid the production of ^13^CO_2_ was detected (mean with SEM ribbon) (n = 3). A) Rate of ^13^CO_2_ production, plotted in nmol/minute. B) Cumulative production of ^13^CO_2_ (mean with individual subject lines plotted). C) The anatomical representation of the location of octanoic acid oxidation in a murine model. D-G) Intraperitoneal injection of saline or metoclopramide 20 minutes before oral gavage of [^13^C] octanoic acid. D) Rate of ^13^CO_2_ production (mean with SEM ribbon, n = 4). E) Cumulative production of ^13^CO_2_. F) The percentage of total tracer oxidized (mean with individual subject lines plotted) during the 200-minute monitoring period. G) The time at which 50 percent of the tracer was oxidized (T_50%_). Tracer 50-percent oxidation time was defined as the time at which half of the total tracer oxidized during the course of the assay, was oxidized. Significant differences were found in the 50-percent oxidation times between the saline and MCP treatment groups. Statistical significance determined by two-tailed unpaired t-test; p < 0.01 (**).

### Mannitol for monitoring oral-cecal transit time (OCTT)

We assessed the kinetics of ^13^C-mannitol tracer oxidation at two sub-milligram doses by measuring rates of ^13^CO_2_ production. We aimed to characterize both the magnitude and temporal profile of ^13^CO_2_ appearance. Notably, ^13^CO_2_ generated from mannitol appeared later than that produced from octanoic acid (**Figure 2A**). Despite administering two different starting doses of mannitol, approximately 15% of the tracer from both groups was oxidized within 6 hours (**Figure 2B**). As expected, the higher mannitol dose resulted in greater total ^13^CO_2_ output (**Figure 2C**). However, no significant difference in T_50%_ (time to 50% oxidation) was observed between the two doses (**Figure 2D**).

**Figure 2:**
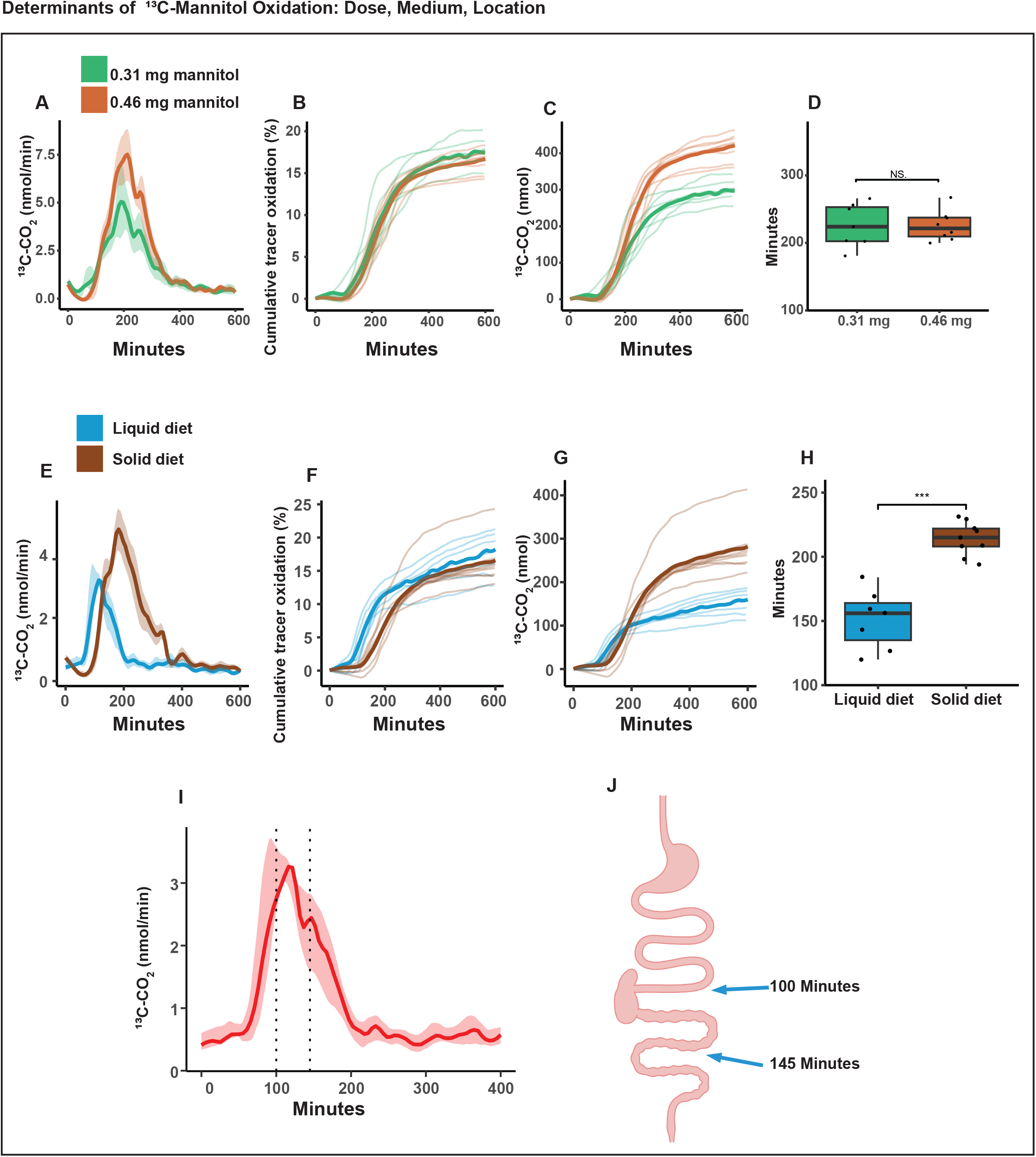
[^13^C]-mannitol breath test in mice. Mice voluntarily ate a [^13^C]-mannitol labeled test meal and the production of ^13^CO_2_ was detected (mean with SEM ribbon) (n = 7-8), plotted by dose. A) Rate of ^13^CO_2_ production, plotted in nmol/minute. B) Cumulative production of ^13^CO_2_ (mean with individual subject lines plotted). C) Cumulative production of ^13^CO_2_. D) T_50%_, in minutes. E-H) [^13^C]-mannitol was gavaged or voluntarily eaten in a test meal and the production of ^13^CO_2_ was detected (mean with SEM ribbon) (N = 7,9). E) ^13^CO_2_ detected (mean with SEM ribbon) after consumption of [^13^C]-mannitol incorporated into peanut butter or after oral gavage of [^13^C]-mannitol (n = 8), grouped by administration strategy. F) The percentage of administered mannitol that was oxidized to ^13^CO_2_ during the course of the assay. G) The total amount of tracer oxidized (mean with individual subject lines plotted) during the course of the assay. H) T_50%_. A statistically significant difference was found between administration strategies. I) ^13^CO_2_ detected (mean with SEM ribbon) after oral gavage of [^13^C]-mannitol (n = 7) J) 100 and 145 minutes after gavage a solution of FD&C1 blue dye, mice were sacrificed and the location of the dye front in the gastrointestinal tract was located. J) graphical representation of the location of the dye after oral gavage of dye. Statistical significance determined by two-tailed unpaired t-test; p ≥ 0.05 (NS), p < 0.001 (***).

Mannitol is widely used as a sugar substitute in processed foods and drinks and is generally well tolerated, indicating its suitability for administration in both food and water. To determine whether the route of administration influences tracer kinetics, we compared oral gavage of mannitol dissolved in water to voluntary consumption mixed with palatable food. For food consumption, we designed the experiment to introduce the tracer to mice with ad libitum access to food and water, as short-term fasting can significantly increase physical activity and metabolic rate in mice (11). Trace amounts of mannitol were mixed with palatable food or administered by oral gavage in water. We observe that liquid gavage of ^13^C-mannitol produced a faster time to peak oxidation when compared to food consumption (**Figure 2E**). Ultimately, the total fraction of oxidized tracer (∼15%) was similar between administration routes (**Figure 2F**). The accelerated kinetics observed with gavage were also apparent in a reduced T_50%_ in this group (**Figure 2H**).

To further explore the sites of mannitol absorption, we tracked gastrointestinal transit using a dye administered by oral gavage (**Figure 2I**). In our tracer studies, maximal ^13^CO_2_ production occurred at 120 minutes post-administration. Accordingly, mice given the dye were sacrificed at 100 minutes and 145 minutes. At 100-minutes, the dye had passed into the small intestine but not the cecum, while at 145 minutes, it had passed the cecum and into the large intestine. These findings indicate that peak ^13^CO_2_ production temporally coincides with tracer arrival in the cecum (**Figure 2J**).

## Microbial Metabolism

While previous studies have shown that mannitol is not absorbed by mammals, we observed that approximately 20% of the administered mannitol tracer was converted to CO_2_. To determine if this oxidation was due to microbial metabolism, we compared tracer oxidation in conventional mice and in mice depleted of commensal microbes using antibiotics, following administration of either ^13^C-mannitol or ^13^C-glucose. Oxidation of ^13^C-glucose was similar in both groups (**Figure 3A–B**). In contrast, antibiotic-treated mice exhibited no detectable ^13^CO_2_ production following ^13^C-mannitol administration (**Figure 3C–D**), indicating that mannitol oxidation depends on the presence of intestinal microbes.

**Figure 3:**
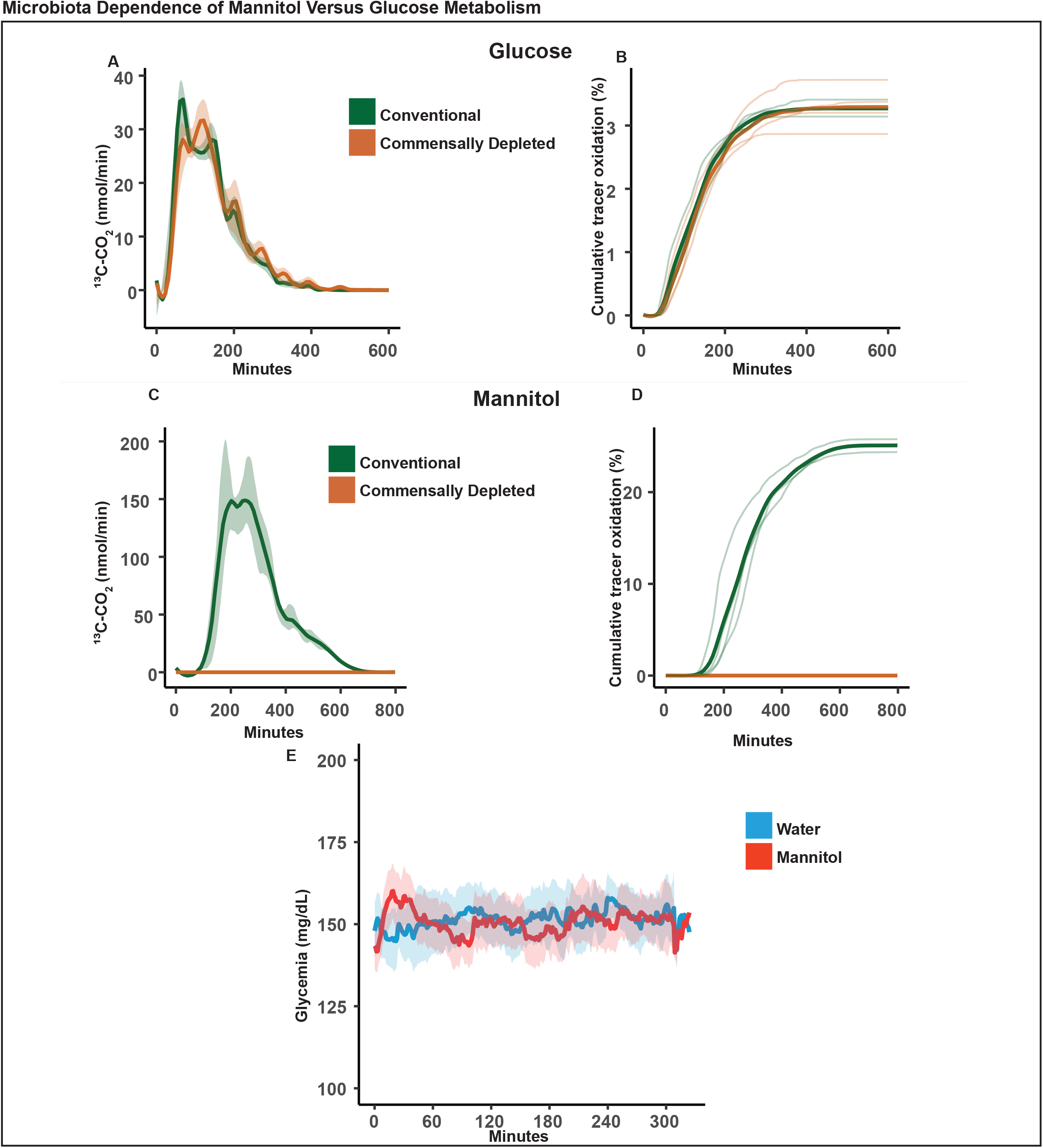
differential [^13^C]-mannitol oxidation in conventional and commensally depleted mice. Mice were gavaged a solution containing [^13^C]-mannitol, [^13^C]-glucose, or water and the production of ^13^CO_2_ was detected (n = 4). A) ^13^CO_2_ detected (mean with SEM ribbon) after oral gavage of [^13^C]-mannitol (N = 4). B) percent of administered mannitol that was oxidized to ^13^CO_2_ during the course of the assay. C) rate of ^13^CO_2_ exhalation (mean with SEM ribbon) after oral gavage of [^13^C] glucose (N=4). D) percent of administered glucose that was oxidized to ^13^CO_2_ during the course of the assay. E) Glycemia measured by implanted continuous glucose monitor after oral gavage of unlabeled-mannitol (N = 3).

Consistent with the literature, mannitol, like other sugar alcohols, is reported to neither increase nor decrease blood glucose, supporting their use as sweeteners for individuals with diabetes (12). Using continuous glucose monitoring, we observed that oral administration of mannitol at a standard glucose tolerance test dose (2 mg/kg) did not raise blood glucose levels over a period corresponding to peak ^13^CO_2_ production (**Figure 3E**).

### Functional measurements: Loperamide

To evaluate the utility of non-invasive OCTT measurements, we performed several functional studies. In the first, we assessed the effect of loperamide, an FDA-approved agent known to reduce gut peristalsis on OCTT (12). Administration of loperamide resulted in a right-shift in ^13^CO_2_ production curves without affecting overall change in tracer absorption (**Figure 4A-B**). This corresponds to a 30-minute delay in time to T_50%_ (**Figure 4C**). In a separate cohort, we tracked the transit of dye through the GI tract following loperamide treatment. In control mice, the dye had reached the colon by 145 minutes, whereas in loperamide-treated mice the dye remained proximal to the cecum (**Figure 4D**).

**Figure 4:**
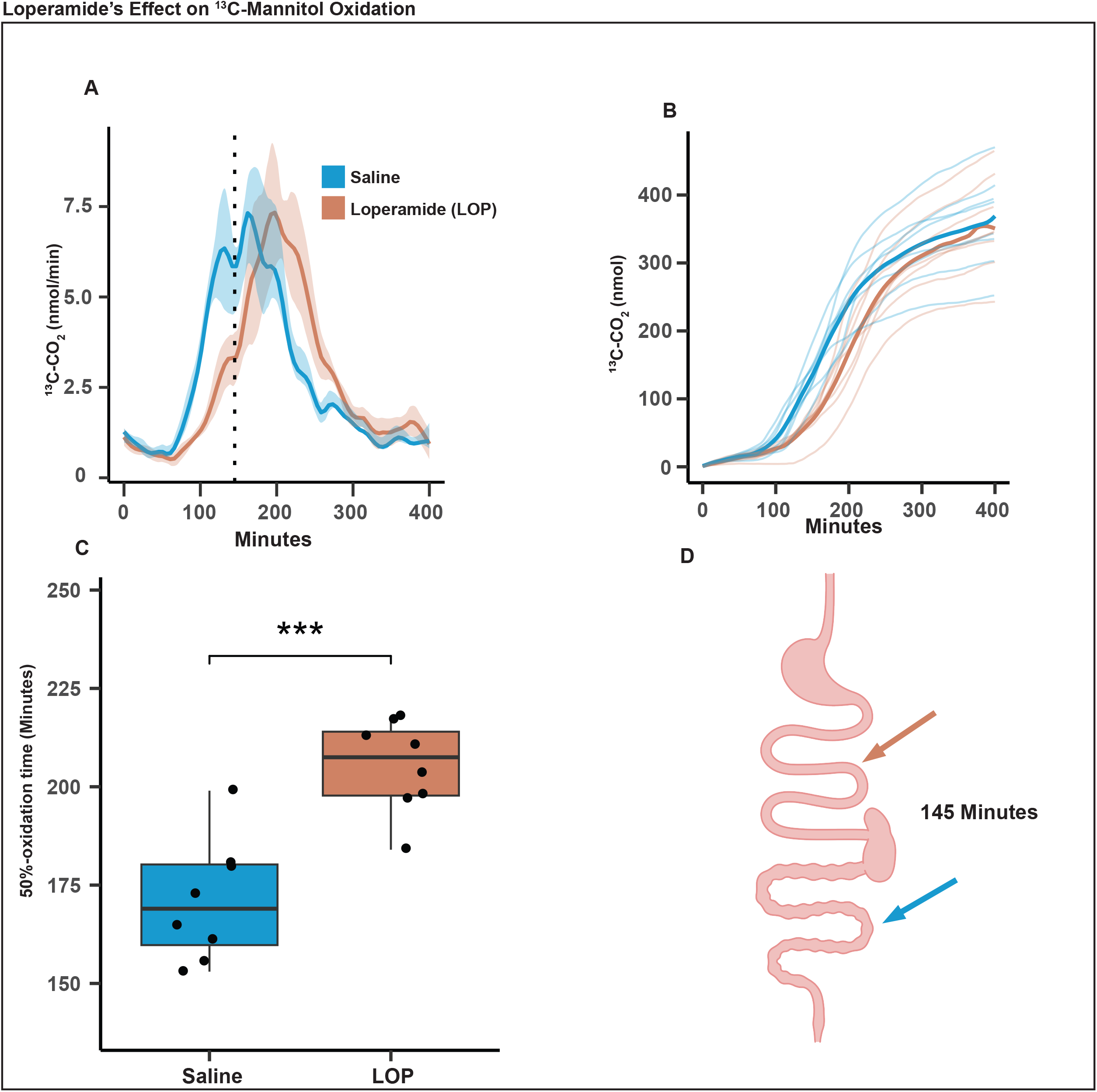
loperamide induced slow-down of mouse gut motility. Mice were treated with loperamide 1-hour prior to gavage of [^13^C]-mannitol, production of ^13^CO_2_ was detected (n = 4). A) Rate of ^13^CO_2_ production, plotted in nmol/minute. B) Cumulative production of ^13^CO_2_ (mean with individual subject lines plotted). C-D) 1-hour after loperamide treatment, 145 minutes after gavage a solution of FD&C1 blue dye, mice were sacrificed and the location of the dye front in the gastrointestinal tract was located. C) T_50%_, a significant increase in T_50%_ was found in the loperamide-treated group relative to control. D) Mice were injected with saline or loperamide, and 1 hour later gavaged with FD&C1 blue dye. After 145 minutes, mice were sacrificed and the position of the dye front in the gastrointestinal tract was determined. graphical representation of the location of the dye front after GI-tract removal of mice gavaged with dye. Statistical significance determined by one-tailed unpaired t-test; p < 0.001 (***).

### Functional measurements: Semaglutide

Finally, we assessed the effect of the GLP1 receptor agonist (GLP1Ra) semaglutide on OCTT which is known to increase gut transit time in humans. In high-fat diet (HFD) fed mice, daily semaglutide treatment resulted in ∼10% reduction in body weight (**Figure 5A**). Semaglutide strongly delayed OCTT and produced a right-shift in the ^13^CO_2_ production curve (**Figure 5B**). Semaglutide treatment reduced overall tracer absorption and delayed T_50%_ times by approximately 100 minutes, a two-thirds increase relative to controls (**Figure 5C-D**). Dye motility assays confirmed this delay with semaglutide significantly slowing dye progression through the GI tract (**Figure 5E**). Energy expenditure and total food consumption were significantly reduced immediately after the start of semaglutide treatment in comparison with PBS treated mice (**Figure 5H-5K**). Consistent with decreased food intake, respiratory exchange ratio (RER) was significantly decreased immediately after start of treatment, and there was a corresponding rapid increase in RER during the washout of semaglutide, with increased food intake and initiation of de novo lipogenesis.

**Figure 5:**
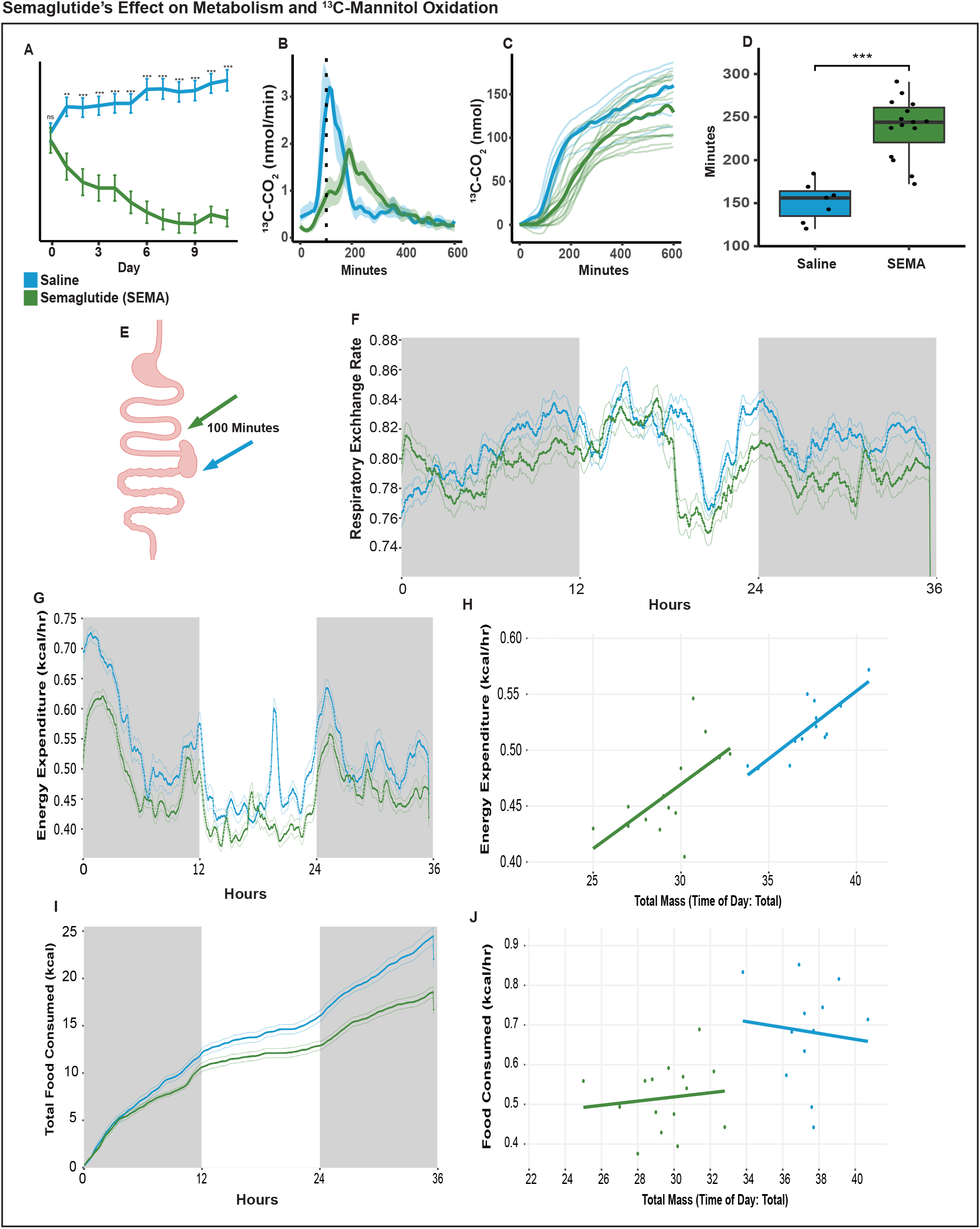
semaglutide induced slow-down of mouse gut motility. Mice were treated with semaglutide (0.06 mg/kg/day) for 7 days prior to gavage of [^13^C]-mannitol, production of ^13^CO_2_ was detected (n = 7,15). A) body weights of mice acutely treated with semaglutide over a 5 day period, with day 0 being baseline (mean with SEM bar, N = 7, 8). B) Rate of ^13^CO_2_ production, plotted in nmol/minute, The dotted line represents the 100-minute mark during the assay. C) Cumulative production of ^13^CO_2_ (mean with individual subject lines plotted). D) T_50%_, a significant increase in T_50%_ was found in the semaglutide-treated group relative to control. E) After daily injection of saline or semaglutide for 7 days, mice were gavaged with FD&C1 blue dye. After 100 minutes, mice were sacrificed and the position of the dye front in the gastrointestinal tract was determined. graphical representation of the location of the dye front after GI-tract removal of mice gavaged with dye. F) Respiratory Exchange Ratio (RER) of mice treated with either semaglutide (0.06 mg/kg/day) or PBS for 9 days followed by 36 hours of observation G-H) Energy expenditure (EE) over the 36 hour window with either semaglutide or PBS treatment and linear regression plot showing the relationship between body weight. A best fit line is shown for both groups. I-J) Food intake over the 36 hour window with either semaglutide or PBS treatment and linear regression plot showing the relationship between body weight and food intake. A best fit line is shown for both groups. Statistical significance determined by two-tailed unpaired t-test; p ≥ 0.05 (NS), p < 0.01 (**), p < 0.001 (***).

## Discussion

Here, we establish a non-invasive, high-resolution method to quantify OCTT in mice by monitoring microbially mediated oxidation of ^13^C-labeled mannitol to ^13^CO_2_. This approach offers several critical advantages over existing methods, including the ability to perform continuous, longitudinal measurements in freely moving, unanesthetized mice without the need for restraint, terminal procedures or radioisotopes. The ability to monitor stable isotope oxidation via breath in real time yields highly granular data on gastrointestinal transit dynamics, with applications for preclinical studies assessing genetic, dietary, microbial, and pharmacological influences on gut motility.

We first validated our system by replicating the well-characterized kinetics of gastric emptying via ^13^C-octanoic acid oxidation. Our data align with prior reports on the utility of the ^13^C-octanoic acid breath test in humans and rodents, supporting the reliability of our multiplexed detection system. Notably, our system detected accelerated gastric emptying with metoclopramide, confirming the assay’s sensitivity to pharmacologically induced changes in motility.

Building on this validation, we identified ^13^C-mannitol as a novel tracer that enables OCTT assessment. Unlike octanoic acid, mannitol is not absorbed by the host and requires microbial metabolism. Our data show that microbial depletion via antibiotics abolishes mannitol oxidation, confirming that the observed ^13^CO_2_ reflects microbial activity rather than host metabolism. These findings are consistent with clinical observations that mannitol does not elevate blood glucose and is a largely non-metabolizable sugar alcohol by mammalian tissues. Our studies do not identify which microbial taxa are capable of mannitol metabolism, an important but unanswered question.

We further demonstrated the functional relevance of this method by pharmacologically altering gut motility. Loperamide treatment delayed the appearance of ^13^CO_2_, consistent with delayed transit to the cecum, as corroborated by dye motility studies. Similarly, semaglutide—a GLP-1 receptor agonist known to slow gastric emptying and intestinal transit—caused a pronounced delay in OCTT in mice on a high-fat diet. These pharmacological challenges validate the sensitivity of our assay to physiologically meaningful changes in gut motility.

Importantly, the route of tracer administration—gavage versus voluntary consumption—altered the kinetics of mannitol oxidation without affecting the total amount oxidized. This flexibility allows the method to be adapted to a variety of experimental designs, including ad libitum administration strategies that minimize stress-related artifacts. It also enables the exploration of how different dietary compositions—such as high-fat, high-protein, high-carbohydrate, or fiber-rich diets—affect gut motility and transit dynamics.

Our approach addresses several limitations inherent to current OCTT methodologies. Terminal methods, while precise, require animal sacrifice at multiple timepoints and are thus resource-intensive and cross-sectional by nature. Other non-terminal approaches, like Carmine red assays, suffer from limited precision and the confounding effects of repeated handling. Breath-based stable isotope analysis circumvents these issues, offering continuous, non-invasive data collection that minimizes stress and variability.

There are some limitations to consider. While mannitol oxidation provides an approximate marker of cecal arrival, its reliance on microbial metabolism could introduce variability in microbiota-depleted or dysbiotic animals. The extent of oxidation may permit a non-invasive measurement of microbiota competent for oxidizing mannitol. However, this inherent reliance on microbial metabolism may complicate interpretation in models expecting major changes to the gut microbiota unless microbial activity is explicitly measured or controlled. Moreover, while we demonstrated equivalency in total oxidation between doses, higher resolution studies may be needed to fully understand dose-dependent kinetics, especially under pathological conditions.

Mannitol is administered clinically for several indications. Mannitol can be used as an osmotic diuretic to reduce acute renal failure or to reduce intracranial pressure when given by intravenous routes (13, 14). Administration of high doses of mannitol in water (e.g. 100 g) will have a laxative effect in bowel preparations (15). Labeled mannitol in trace doses is unlikely to cause significant side effects in patients or in rodents.

In summary, our work establishes a scalable, non-invasive method to quantify OCTT with high temporal resolution in mice, opening new avenues to investigate and visualize gut motility and host-microbe interactions. This method enhances the translational potential of murine models by reducing animal use, limiting experimental confounds, and enabling dynamic assessment of gastrointestinal function in response to genetic, environmental, and pharmacologic factors.

## Methods

### Animals

All animal experiments were approved by the Beth Israel Deaconess Medical Center IACUC. Wildtype, male C57BL/6 mice (Jackson Laboratory, Strain No. 000664) were used in all animal experiments. Mice were maintained at 12 h/ 12 h light/ dark cycles 0800/1600, 22 ± 2 °C room temperature and 30%–70% humidity with ad libitum access to food and water, in individually ventilated cages. Cages and bedding were changed once every two weeks, and mice were monitored regularly for their health status by animal technicians with the support of veterinary care and remained free of any adventitious infections for the entire duration of this study. Mice were fed a standard chow diet (13% kcal fat, LabDiet, no. 5008) or a high fat diet (HFD, 60% kcal fat, Research Diets, no. D12492i) for the indicated durations. All experiments were performed in adult mice (12 to 24 weeks of age). For semaglutide diet comparison studies, mice were placed on HFD 4 weeks prior to the start of the studies.

### Peanut butter acclimation

While octanoic acid tracer incorporation into cooked egg yolks has traditionally been used with patients and with rats, we found inconsistent consumption of this test meal in C57Bl/6J mice (4, 16). Therefore, to promote the voluntary consumption of our tracer compounds, peanut butter was selected as a palatable alternative. Mice were acclimated to peanut butter with either unlabeled mannitol or octanoic acid in a ceramic dish for 30 minutes on 3 successive days.

### ^13^C-octanoic acid gastric emptying assay

We used 1-^13^C-labeled octanoic acid (Cambridge Isotope Laboratories, CLM-293) to validate gastric emptying measurements. 160 µL of liquid octanoic acid was diluted in 1 mL DMSO, diluted in sterile water to 5.4 mg/mL, and 160 µL of this solution was mixed into 5 g of peanut butter (Jif, WB Mason, SMU5150024114) warmed to 60 °C, stirred, then cooled to room temperature. Mice were provided a 100-mg portion of the ^13^C-octanoic acid– enriched peanut butter in a clean ceramic dish and monitored to confirm full consumption within 30 minutes. Mice that did not fully consume the dose were excluded. Mice had free access to food and water for the duration of the experiment. Exhaled breath was sampled continuously for up to 200 minutes using an ABB Off-Axis Integrated Cavity Output Spectroscopy (OA-ICOS) gas analyzer connected to a Promethion indirect calorimetry system (Sable Systems). Data were exported from Promethion Live (Sable Systems, v24.0.12), converted to CalR-compatible files, and analyzed using RStudio (version 2024.12.1 Build 563, “Kousa Dogwood” Release) with R version 4.4.1. For pharmacological validation studies, a subset of mice received 10 mg/kg metoclopramide, intraperitoneally (Sigma Aldrich, M0763) or saline 20 minutes before administration of a 200-µL oral gavage of ^13^C-octanoic acid in water.

### Antibiotic Induced Commensal Depletion

To deplete commensal bacteria, mice were provided with an antibiotic cocktail in their drinking water. Antibiotics were dissolved in autoclaved, filtered facility water, adjusted to a neutral pH (7.4), and filter-sterilized prior to use. The medicated water was provided in light-protected bottles and replaced every 5–7 days to ensure stability and minimize microbial or fungal contamination. The antibiotic cocktail consisted of neomycin trisulfate (1 g/L; Sigma-Aldrich, N5285-25G), metronidazole (0.25 g/L; Patterson Vet Generics, 78912046), vancomycin hydrochloride (0.5 g/L; Sigma-Aldrich, V8138-1G), and ampicillin trihydrate (1 g/L; Sigma-Aldrich, A1593-25G). Neomycin and ampicillin were prepared from powder, while metronidazole was added as a liquid injectable formulation (500 mg/100 mL), at a dose of 0.25 g/L.

### ^13^C-mannitol oral-cecal transit time assay

Oral-cecal transit time was assessed using ^13^C-labeled mannitol (Cambridge Isotope Laboratories, CLM-6733). The mannitol was received as a powder and dissolved to 100 mg/mL in sterile water to prepare it for administration. For voluntary feeding studies, 160 µL of stock was mixed into 5 g of peanut butter and cooled after heating to 60 °C. For gavage, the solution was diluted to 0.8 mg/mL in sterile water. Mice received either a 100-mg portion of the prepared peanut butter or a 200-µL oral gavage of ^13^C-mannitol. Mice had free access to food and water for the duration of the experiment. Breath samples were collected continuously for up to 6 hours with the Promethion and ABB OA-ICOS. Data processing and analysis were performed in RStudio following conversion to CalR-compatible format. In separate experiments, FD&C Blue No. 1 dye (200 µL of dyed water) was administered by gavage to map the dye front at specific timepoints (100 or 145 minutes) post-gavage. After sacrifice, the gastrointestinal tract was examined and photographed.

### Functional modulation of oral-cecal transit time

To test whether the system could detect pharmacologically induced changes in OCTT, mice received agents known to alter gastrointestinal motility. Loperamide hydrochloride (1 mg/kg; Sigma, L4762) was prepared in 100% ethanol and saline and administered intraperitoneally 1 hour before ^13^C-mannitol gavage. Semaglutide sodium salt (0.06 mg/kg/day; BOC Sciences, B0084-007194) was prepared in saline and administered subcutaneously daily for one week before and during ^13^C-mannitol testing in mice on a 60% high-fat diet (Research Diets, D12492i). Breath sampling and data analysis were conducted as previously described. For dye transit studies, was administered by gavage to mice, and their gastrointestinal tracts were examined at predetermined time points to assess the location of the dye front. Measurements of metabolic rate were collected in the Promethion High Definition Multiplexed Respirometry System (Sable Systems) for the semaglutide study. Following a 7 day injection schedule, mice were monitored with this system for 36 hours. Measurement of VO_2_, VCO_2_, RER, locomotor and ambulatory activity, food intake was done at 23 ± 0.1°C. Energy expenditure was calculated with the Weir equation (Weir, 1949).

### Semaglutide Metabolic Parameters Assay

To show the effect of semaglutide and the rapid changes in metabolism after semaglutide treatment, 16 week old male mice on a HFD were injected daily with either PBS or Semaglutide sodium salt subcutaneously for 7 days. Measurements of metabolic rate were collected in the Promethion High Definition Multiplexed Respirometry System (Sable Systems) following this injection schedule, mice were monitored within this system for 36 hours while injections continued to occur daily. Measurement of VO_2_, VCO_2_, RER, locomotor and ambulatory activity, food intake were done at 23 ± 0.1°C. On day 9, mice were gavaged with ^13^C-labeled mannitol, prepared as previously described, to measure changes in OCTT. 3 out of 16 mice in the PBS group were excluded for poor quality data. 2 out of 16 mice in the semaglutide group were excluded for poor quality data.

## Data Analysis

Data were processed and analyzed using R (version 2024.12.1; RStudio version 4.4.1). The following pipeline was used:

### Data Preparation

Raw data were exported from CalR and imported into R. Feeding start times were aligned across subjects using a pre-recorded offset. An adjusted experimental time variable was calculated by subtracting each subject’s offset from their recorded timepoints. Baseline Correction: Baseline CO_2_ production and ^13^C-enrichment (C13) were calculated over a defined pre-treatment window for each subject. Baseline-subtracted (adjusted) values were computed for both variables. Filtering and Quality Control: Data were filtered to focus on the experimental period of interest (e.g., 0–600 minutes post-feeding). Visual inspection of C13 × vCO_2_ plots for individual subjects was used to identify and exclude data artifacts or outliers.

### Normalization and Calculations

VCO_2_ was converted to μmol/min and subsequently to nmol/min. Cumulative sums of CO_2_ production and tracer oxidation were calculated. Total tracer availability (in μmol) was determined based on dosing information. The percentage of tracer oxidized over time was calculated per subject.

Data Cleaning: Super-negative ratios during cage adjustments (such as during tracer administration) were removed prior to plotting and statistical analysis. Data Aggregation: Separate datasets for each condition (e.g., low vs. high mannitol dose) were processed and combined for comparative analysis.

### Statistical Analysis

The time to 50% tracer oxidation was calculated per subject. Group comparisons were performed using Welch’s t-test (unequal variances). Sample sizes, means, standard deviations, standard errors, t-statistics, degrees of freedom, p-values (two-tailed), 95% confidence intervals, pooled standard deviations, and Cohen’s d effect sizes were computed. Boxplots were generated to visualize group differences in 50% oxidation times, annotated with significance markers.

## Acknowledgements

All of the graphical abstract and parts figures 1, 2, 4, and 5 were created with BioRender.com. We would like to thank Ellen Vaynberg, Brad Fortunato, Subhash Kulkarni, Brad Lowell, Angela Kim, Marshall McCue for support and helpful discussions.

---

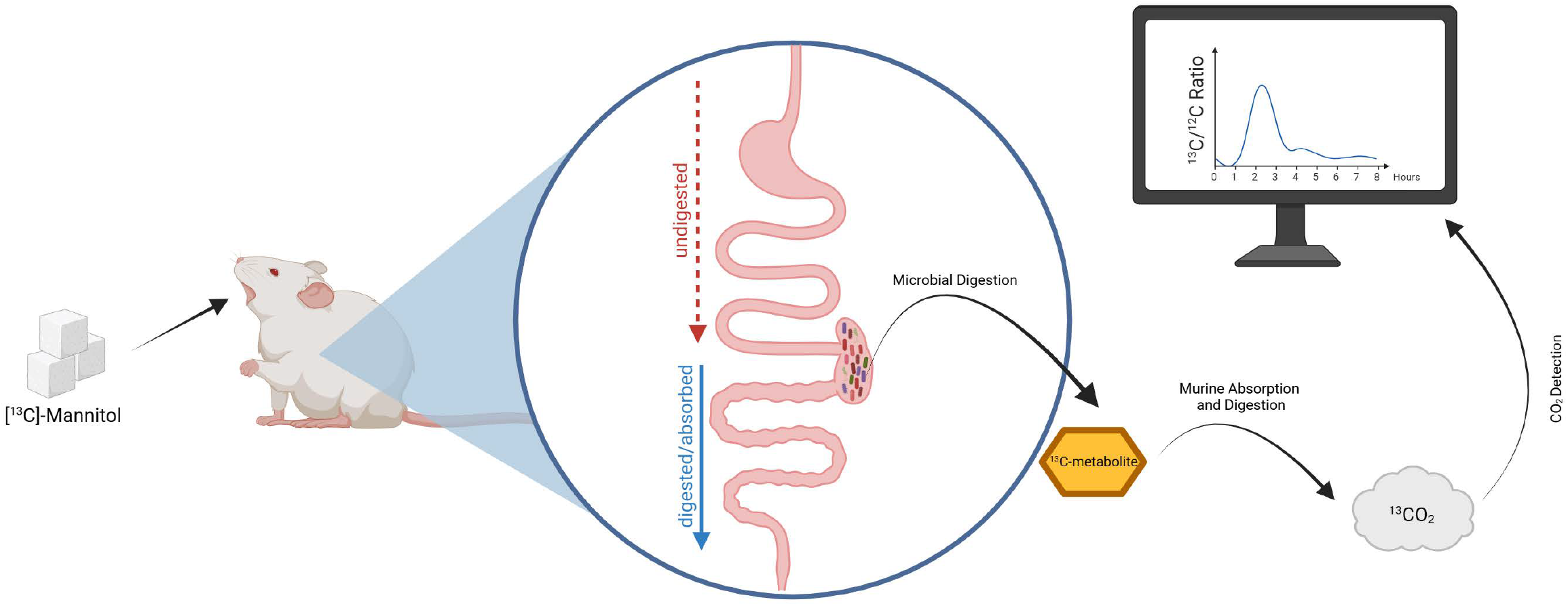

